# Defining the core essential genome of *Pseudomonas aeruginosa*

**DOI:** 10.1101/396689

**Authors:** Bradley E. Poulsen, Rui Yang, Anne E. Clatworthy, Tiantian White, Sarah J. Osmulski, Li Li, Cristina Penaranda, Eric S. Lander, Noam Shoresh, Deborah T. Hung

## Abstract

Genomics offered the promise of transforming antibiotic discovery by revealing many new essential genes as good targets, but the results fell short of the promise. It is becoming clear that a major limitation was that essential genes for a bacterial species were often defined based on a single or limited number of strains grown under a single or limited number of *in vitro* laboratory conditions. In fact, the essentiality of a gene can depend on both genetic background and growth condition. We thus developed a strategy for more rigorously defining the core essential genome of a bacterial species by studying many pathogen strains and growth conditions. We assessed how many strains must be examined to converge on a set of core essential genes for a species. We used transposon insertion sequencing (Tn-Seq) to define essential genes in nine strains of *Pseudomonas aeruginosa* on five different media and developed a novel statistical model, *FiTnEss*, to classify genes as essential versus non-essential across all strain-media combinations. We defined a set of 321 core essential genes, representing 6.6% of the genome. We determined that analysis of 4 strains was typically sufficient in *P. aeruginosa* to converge on a set of core essential genes likely to be essential across the species across a wide range of conditions relevant to *in vivo* infection, and thus to represent attractive targets for novel drug discovery.

---

The sequencing of the first bacterial genome in 1995 (1), offered the promise of revolutionizing antibiotic discovery by revealing the breadth of genes that could be mined for antibiotic targets and paved the way for genome-wide genetic screens to identify essential genes in a given bacterial species and chemical screens to find new antibiotics inhibiting these essential targets. However, the experiences of two major pharmaceutical companies in the late 1990s to early 2000s suggest that this promise was not fulfilled (2, 3). While several factors contributed to the disappointing yield of new antibiotic candidates, one important factor was that inhibitors of supposedly essential targets often failed to have good activity against all pathogen strains or to clear infections.

In retrospect, it is becoming clear that the criteria used at the time for declaring a gene to be essential within a species were not sufficiently rigorous, with essentiality often defined based on the effect of inactivating a gene in a single strain of a pathogen species under a single *in vitro*, laboratory growth condition. We now recognize that whether a gene is essential may depend on both genetic background (i.e, the strain in which it resides) (2) and growth conditions (*i.e.*, conditional essentiality) (4, 5). Thus, given the diversity of bacterial genomes even within a species, genes essential in a single strain need not be essential in all strains of a given species. And, given the variable environments encountered by bacterial pathogens in lab media and different infection types (*i.e*., blood, urine, lung, abscess infections), genes essential in artificial laboratory growth conditions need not be essential during human infection. Focusing on ‘core essential genes’ – by which we mean genes that are essential across virtually all strains of a pathogen species and all relevant growth conditions – would likely increase the success of antibiotic discovery. We therefore sought to develop a robust paradigm for defining the core essential genes of a bacterial species.

We focused on *Pseudomonas aeruginosa*, a clinically significant pathogen that is a major cause of bacteremia, pulmonary, and urinary tract infections, with high mortality rates (6-8), and for which there is the greatest need for new antibiotics. Due to its ability to evade current antibiotics or develop resistance, *P. aeruginosa* clinical strains are increasingly resistant to all current antibiotics (9, 10). The World Health Organization has recently classified *P. aeruginosa* as a priority pathogen in need of research investment and new drugs (11). Alarmingly, only 1 in 5 antibacterial drugs succeed in clinical trials (12), and of the 48 potential antibacterials in development as of 2018, only 3 have activity against *P. aeruginosa* with only 1 of these having a new mechanism of action (www.pewtrusts.org/antibiotic-pipeline).

Here we examine two fundamental questions: How accurately can the core essential genome be identified based on essentiality in one strain under one laboratory growth condition? How many strains must be examined to converge on a set of core essential genes that are likely to be essential under conditions relevant for infection and thus may be good drug targets? We addressed this question by using Tn-Seq (also known as TIS, INseq, HITS, TraDIS, (13-17)) to perform genome-wide negative selection studies on libraries of transposon-insertion mutants under different growth conditions, with the distribution of transposon insertions determined by sequencing the pool of strains. Genes that are important for optimal growth under a specific growth condition can be identified, because the corresponding mutants containing disrupting transposon insertions in these genes will be seriously depleted from the pool of all possible mutants. These methods have been applied to the two commonly studied reference lab strains of *P. aeruginosa*, PA14 and PAO1, with varying numbers and identities of essential genes (18). Here, we applied this method to PA14 on Luria-Bertani (LB) media and compared the essential genome determined from this single strain on a single lab-based media to 8 other diverse strains of *P. aeruginosa* under 5 different growth conditions. The strains comprised isolates from various human infections (including pulmonary, urinary, blood, wound and ocular) and one environmentally isolated strain, while the growth conditions comprised three media intended to simulate the conditions of human infection (sputum, serum, urine) and two lab-based media (LB and M9 minimal media). We further developed a novel, simple statistical method, called *FiTnEss* (<u>Fi</u>nding <u>Tn</u>-Seq <u>Ess</u>ential genes), that maps measurements of fitness of individual transposon mutants onto a binary classification of essential or nonessential with user defined levels of stringency. We applied *FiTnEss* to the Tn-Seq data from all strain and media combinations and defined a set of 321 core essential genes, which represent 6.6% of the genome, that constitute a high-priority list of candidate targets for drug discovery against this important pathogen. Finally, we calculated that as few as 4 individual species could be examined in combination to approach a plateau of core essential genes across a given species.

## Results

### Transposon mutagenesis, sequencing, and mapping of transposon insertions

We chose strains from a collection of 130 clinical *P. aeruginosa* isolates obtained from various sources (see Methods). After performing whole genome sequencing of the collection, mapping the isolates to a phylogenetic tree formed by 2560 *P. aeruginosa* genomes in NCBI, and testing a subset for their ability to be efficiently mutagenized by the Himar1-derived transposon MAR2xT7 (19-21), we focused on nine strains that represented five different infection types (blood, urine, respiratory, ocular and wound), with each strain representing a different branch of the dendrogram (NCBI; Fig. 1A). The genomes of these 9 strains varied from 6.34 to 7.15 Mbp.

**Figure 1.**
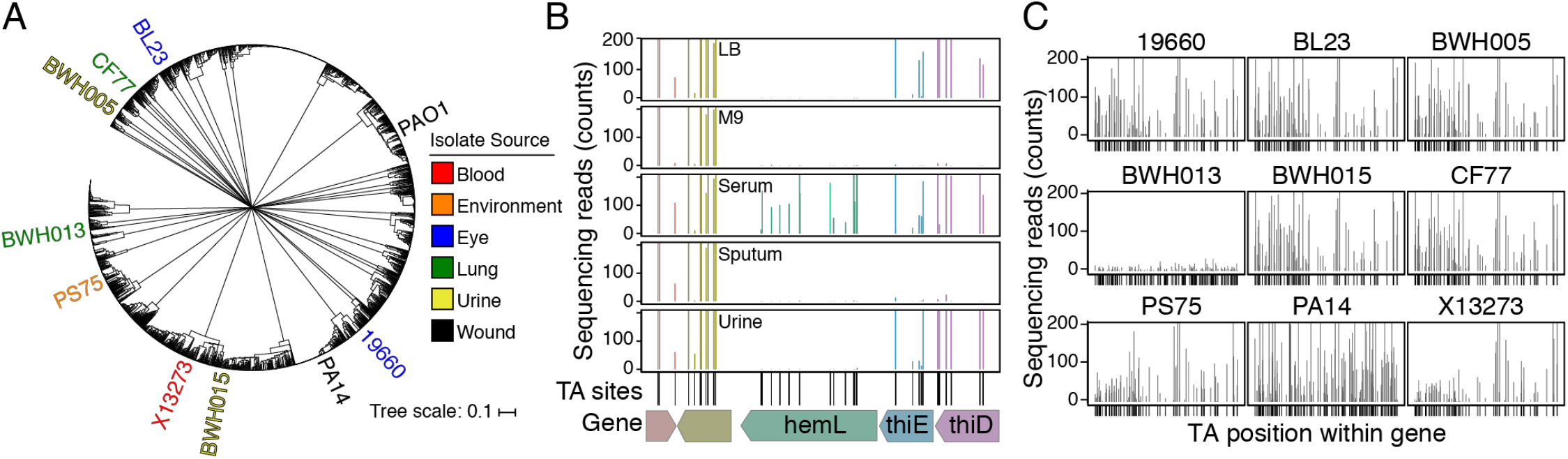
Tn-seq of *P. aeruginosa* clinical isolates. A. Phylogenetic dendrogram of 2560 *P. aeruginosa* genomes (NCBI); PAO1 and strains selected for mutagenesis are indicated. B. The variable sequencing reads which map to TA sites in an exemplary region of five genes in strain PA14 (including *hemL, thiE* and *thiD*) under different growth conditions highlights the conditional essentiality of these genes. C. Normalized read counts mapping to the *pilY1* gene in all nine strains in LB medium demonstrates the variable essentiality of *pilY1* in different strains as an example of the genomic heterogeneity of *P. aeruginosa* isolates.

We constructed transposon libraries by performing tripartite matings of these 9 *P. aeruginosa* strains with *E. coli* donor strain SM10 carrying an episomal MAR2xT7 transposon (20) and *E. coli* strain SM10 carrying an episomal hyperactive transposase that results in efficient integration at the dinucleotide sequence ‘TA’ (Fig. S1) (22). Separating the transposase and transposon increased the efficiency of insertion sequencing and mapping, relative to the more common system of a single plasmid carrying both the transposase and the transposon. We obtained at least 5×10^6^ distinct mutants for each strain from at least two independent conjugations, and selected mutants on each of five different agar media directly to avoid a bottleneck from pre-selecting the libraries on a given medium. To ensure saturating mutagenesis, a total of lx10^6^ mutants were selected on each medium in duplicate, yielding 10-fold more transposon mutants than possible insertion sites. The media types included rich (LB) and minimal (M9) laboratory media, to provide the boundaries (extremes of growth conditions) for essential gene identification, and three media intended to resemble infection site fluids: fetal bovine serum, synthetic cystic fibrosis sputum (23), and urine. We mapped the transposon-insertion sites to the corresponding reference genomes for each strain.

In all, we created 90 Tn-Seq datasets (9 strains grown on 5 media, performed in duplicate), with an average number of mapped reads of approximately 10^7^. Reads at each TA site were highly concordant between replicates, with a mean R^2^ = 0.98 (Dataset S1).

Visual inspection readily identified examples of genes that were variably essential under different growth conditions for a certain strain, illustrating the conditional essentiality of some genes (Fig. 1B). For example, the thiamine synthesis genes *thiD* and *thiE* showed few insertions in M9 minimal media, which lacks thiamine, but an abundance of insertions in rich LB media, indicating their essentiality in M9 but not LB. Variability is also seen for the *hemL* gene under different growth conditions. We similarly saw examples of genes that were variably essential in different strains under the same growth condition. For example, the *pilY1* gene did not tolerate insertions in strain BWH013, but readily tolerated insertions in the other 8 strains, when grown on LB — highlighting the genomic plasticity of *P. aeruginosa* (Fig. 1C).

To optimize our accuracy in calling genes essential or non-essential, we removed from our analysis three classes of TA sites that can lead to technical errors. These classes include (1) non-permissive insertion sites consisting of the sequence (GC)GNTANC(GC), which was recently reported to be intolerant to Himar1 transposon insertions in *Mycobacterium tuberculosis* (24) and which we confirmed is also intolerant in *P. aeruginosa*; (2) non-disruptive terminal insertions within 50 bps of the 5’- and 3’-gene termini (a distance we optimized empirically) which can nevertheless result in the expression of a functional, albeit truncated version of the corresponding gene product (25); and (3) insertion sequences at which genomic sequences flanking a TA site were not unique and could not be accurately mapped (Fig. S2 and methods). In total, we removed 16,499 of 81,328 TA sites (20%) in PA14, which resulted in our inability to assess 150 genes in PA14 (2.5%). The inability to assess the essentiality of genes that contain zero TA sites (35) removed another 185 genes from analysis in PA14. In total we were able to assess the essentiality of 5708 of the 5893 total genes in the PA14 genome (97%). The statistics were similar for the other 8 strains (Table S1).

### *FiTnEss*: a statistical model to identify essential genes

We next sought to perform a comprehensive and quantitative analysis of the 90 Tn-Seq datasets. While various methods exist for analyzing Tn-Seq data (13, 26-28), they differ in their complexity and their stringency for calling a gene as essential. We thus developed a simple model and method (*FiTnEss*, <u>Fi</u>nding <u>Tn</u>-Seq <u>Es</u>sentials) for identifying essential genes from Tn-Seq data that required minimal assumptions and had good predictive power. Importantly, we evaluated essentiality at the level of genes rather than individual TA sites. Across the entire dataset, we found that the average number of reads per TA site for a gene (ng/NTA where ng is total number of reads across the gene and NTA is number of TA sites) falls into a clear bimodal distribution, with presumed essential genes on the left (with a small or zero average ng/NTA) and non-essential genes on the right (ng/NTA > 0) (Fig. S3). However, we found that this average (ng/NTA) varies based on gene length, due to statistical averaging; in contrast, we empirically observed that the read-number from randomly selected, individual TA sites in nonessential genes are similar, regardless of the length of the gene in which they are contained (Fig. S3), suggesting that they are dependent only on mutant fitness. Given these findings, we based *FiTnEss* on modeling the read-number distribution for randomly selected, individual TA sites in clearly non-essential genes (NTA = 10; top 75% of the distribution) and determined the model parameters from the data. We posited that this distribution is geometric with probability (pg) and that 1/pg further follows a lognormal distribution. Thus, requiring only two parameters, the mean and the variance of a distribution, we were able to accurately capture the behavior of all nonessential genes (Fig. S3).

Using the two parameters (determined individually for each dataset), we then constructed a theoretical ‘nonessential’ distribution for different gene sizes for each corresponding dataset, and calculated the probability (p-value) of a given gene coming from this non-essential distribution. In order to vary the stringency with which we called essentiality, we applied two different levels of multiple testing adjustment: one with maximal stringency to offer the highest confidence set of essential genes (family-wise error rate (FWER)) to identify genes with no or very few sequencing reads); and one with high stringency, yet slightly relaxed (false discovery rate (FDR)) to identify genes that are statistically significant yet contain a low number of reads). Genes with an adjusted p-value < 0.05 in both replicates were predicted to be essential (Fig. 2A and SI methods). Virtually all maximal stringency calls are expected to be true essential genes, while among the high stringency set, a small number of false positives is expected.

**Figure 2.**
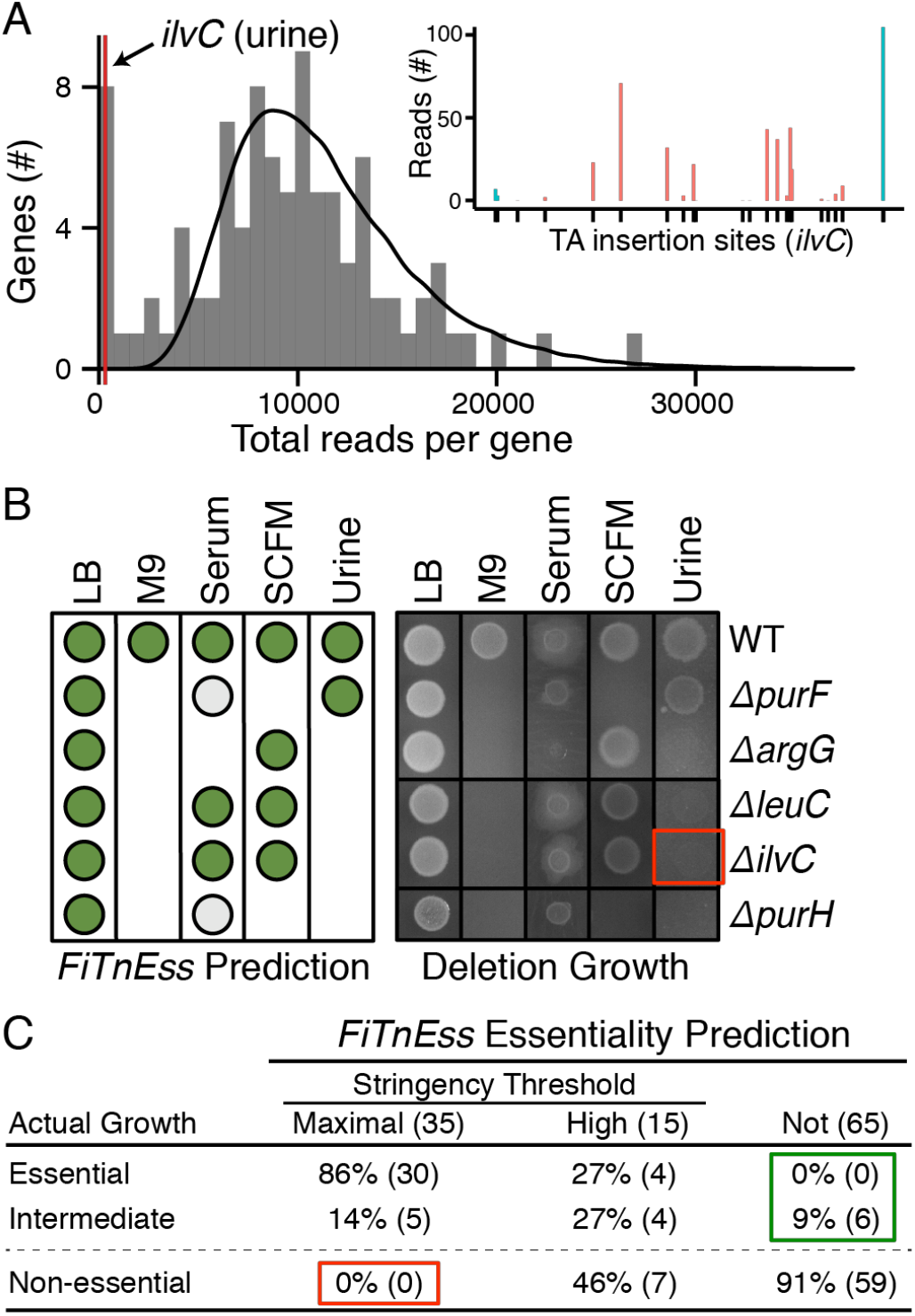
Validation of *FiTnEss* predictions on a set of conditionally essential gene deletions. A. *FiTnEss* prediction for *ilvC* in urine. The grey histogram is the actual distribution of Tn-seq reads for all genes with 18 insertion sites in PA14 grown in urine, the black line is the theoretical distribution calculated by *FiTnEss*, and the red line is where *ilvC* falls at the far left (*i.e*. essential) of the bimodal distribution. Inset: read numbers at useable (red) and removed (blue) TA sites are shown. B. *FiTnEss* essentiality predictions (left; non-essential, high stringency essential, and maximal stringency essential predictions displayed as dark green circles, light green circles, and blank spaces, respectively) of five representative gene deletion mutants from PA14; actual mutant growth mirrored predicted growth on 5 media (right). The red box identifies the absence of growth of Δ*ilvC* (the deletion mutant highlighted in panel A), thus experimentally confirming its essentiality on urine. The full growth profiles of 23 gene deletions can be found in Fig. S4. C. A summary of *FiTnEss* performance based on actual deletion mutant growth profiles. Gene-medium instances are indicated in parentheses; red and green boxes highlight false positive and negative rates, respectively.

### Validating *FiTnEss* using strain PA14

To validate *FiTnEss*’s approach to predicting gene essentiality, we compared its predictions to actual viability and growth measurements for a set of PA14 mutants in which we cleanly deleted particular genes of interest. We created clean deletion mutants corresponding to 20 genes that *FiTnEss* identified as non-essential in LB, but were essential in one or more of the other media, as well as to 3 control genes that were predicted to be non-essential in all media. We determined the positive and negative predictive values of *FiTnEss* by growing the 23 mutants on the same five media as used in the original Tn-Seq experiments, for a total of 115 gene-medium combinations. Mutant strain viability was categorized as essential, intermediate, and non-essential using densitometry (<20%, 20-50%, and >50% relative to WT, respectively; Fig. 2B-C and Fig. S4). Of the 35 combinations predicted to be essential by the maximal stringency criteria, 30 were indeed found to be as essential and 5 were of intermediate growth. Importantly, no strains within this criterion were in fact non-essential. By relaxing the stringency slightly to ‘highly stringent’, 15 additional strain-medium combinations were predicted to be essential, 8 of which were truly essential or of intermediate growth and the remaining 7 were non-essential, corroborating our prediction that some false positives would be expected in this category. Of the 65 combinations predicted to be nonessential, none were found as essential, but 6 instances were found to be of intermediate growth. In this limited dataset, *FiTnEss* had a positive predictive value of 100% by using maximal stringency and 86% using the high stringency predictions (if intermediate genes are classified as essential). The negative predictive value of *FiTnEss* is 91% or 100% (depending on the classification of intermediate genes). Notably, all 3 control strains behaved as predicted, growing on all media, despite our having chosen these control strains with p-values that fall at the boundary drawn to distinguish essential and non-essential genes, further reinforcing the accuracy of this binary classification. All together, these results support *FiTnEss’s* ability to accurately call essential and non-essential genes, and that *FiTnEss’s* stringency can be varied based on user tolerance of false positive versus false negative predictions. Importantly, *FiTnEss* correctly predicted gene essentiality despite the presence of a small number of mapped insertions in the primary Tn-Seq data of some genes, as exemplified in the case of the *ilvC* gene encoding ketol-acid reductoisomerase (Fig. 2A).

### Defining the core genome

We first defined the core genome consisting of 5109 protein-coding genes (genes present in all 9 strains) using the orthogroup clustering software Synerclust (29). The size of the core genome defined by these 9 strains is comparable to what has been previously described for *P. aeruginosa* (5316 total genes, (30)). Of these genes, 4903 were present in single-copy with TA sites that allowed assessment by Tn-Seq; the remaining genes (86 multi-copy genes in which reads could not be accurately mapped due to sequence homology, and 120 small genes that do not have TA sites permissive to transposon insertion) could not be assessed (Dataset S2). The accessory genome within each strain (the genes that are present in the strain but not in all strains) ranged from 655-1369 genes.

### Defining the core essential genome

We then examined the *FiTnEss* predictions for all 90 datasets to identify the core essential genes across the *P. aeruginosa* species (Dataset S3 and Table 1). If one examines only a single strain in a single medium, the number of essential genes varies widely between 354 and 727 genes, even when using the maximal stringency prediction (Table 1). If one examines only genes common to all strains (the core genome), however, the number of essential genes from strain to strain was much more tightly distributed (337 to 386; Fig. S5). In contrast, the number of essential genes in the accessory genome of each strain varied widely from 59 to 478 genes (Fig. S5); interestingly, the number was roughly proportional to genome size (Fig. S5).

When combining all 9 strains across the 5 media and applying maximal stringency, we found that there are only 249 core essential genes (5.1% of the genome). This number is up to three-fold fewer than the number found for a single strain and medium. If we apply the slightly lower standard of high stringency (to allow for the possibility of some false negatives in the data), an additional 72 genes (1.5% of the genome) are included — resulting in 321 genes. We define this set as the core essential genome.

To assess whether the number of core essential genome had reached a plateau, we calculated how the number of essential genes decreases with additional strains (with strains added in 10,000 different random orders) (Fig. 3A). We found that the median across these trajectories typically plateaued after 4 strains, but that 5 strains would ensure 90% of all trajectories reaching a plateau defined as a < 5% false positive rate (Fig. 3B). Beyond the 4 strains, the maximum number of core essential genes declined by only 13 genes. These genes just failed to reach the essential p-value threshold, suggesting that they were false negatives (Dataset S4). If we were to include these 13 genes, the core essential genome would reach 334 genes.

**Figure 3.**
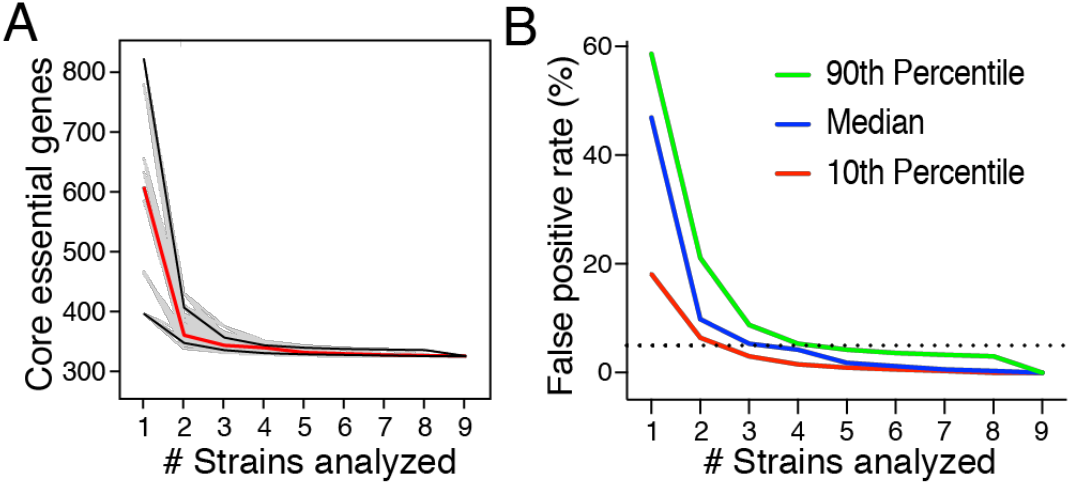
Core essential genome definition plateau. A. 10,000 random calculations (grey) of the trajectory of the number of core essential genes determined upon the sequential introduction of additional strains, up to a total of 9 strains. The 10^th^ and 90^th^ percentiles (black lines) and mean (red line) calculated trajectories of core essential genome sizes are highlighted. B. False positive rate of core essential genes upon the introduction of strains as calculated in A. The dashed line represents a 5% false positive rate, with the median (blue) number of strains to cross this threshold being 4, and the 10^th^ (red) and 90^th^ percentile (green) crossing the threshold at 3 and 5 strains, respectively.

We examined the identities and functions of the core essential genes. Of the 321 core essential genes, 263 correspond to cytosolic proteins, with 132 involved in metabolic pathways (50%) and 119 involved in macromolecular synthesis including DNA replication, transcription or translation (45%). Another 56 correspond to cytoplasmic membrane, periplasmic and outer membrane proteins with the majority involved in cell structure and division, metabolism, or act as transporters/chaperones (13, 12 and 26 genes, respectively). The remaining 12 of the 321 genes are completely uncharacterized (Fig. 4A and Dataset S5).

**Figure 4.**
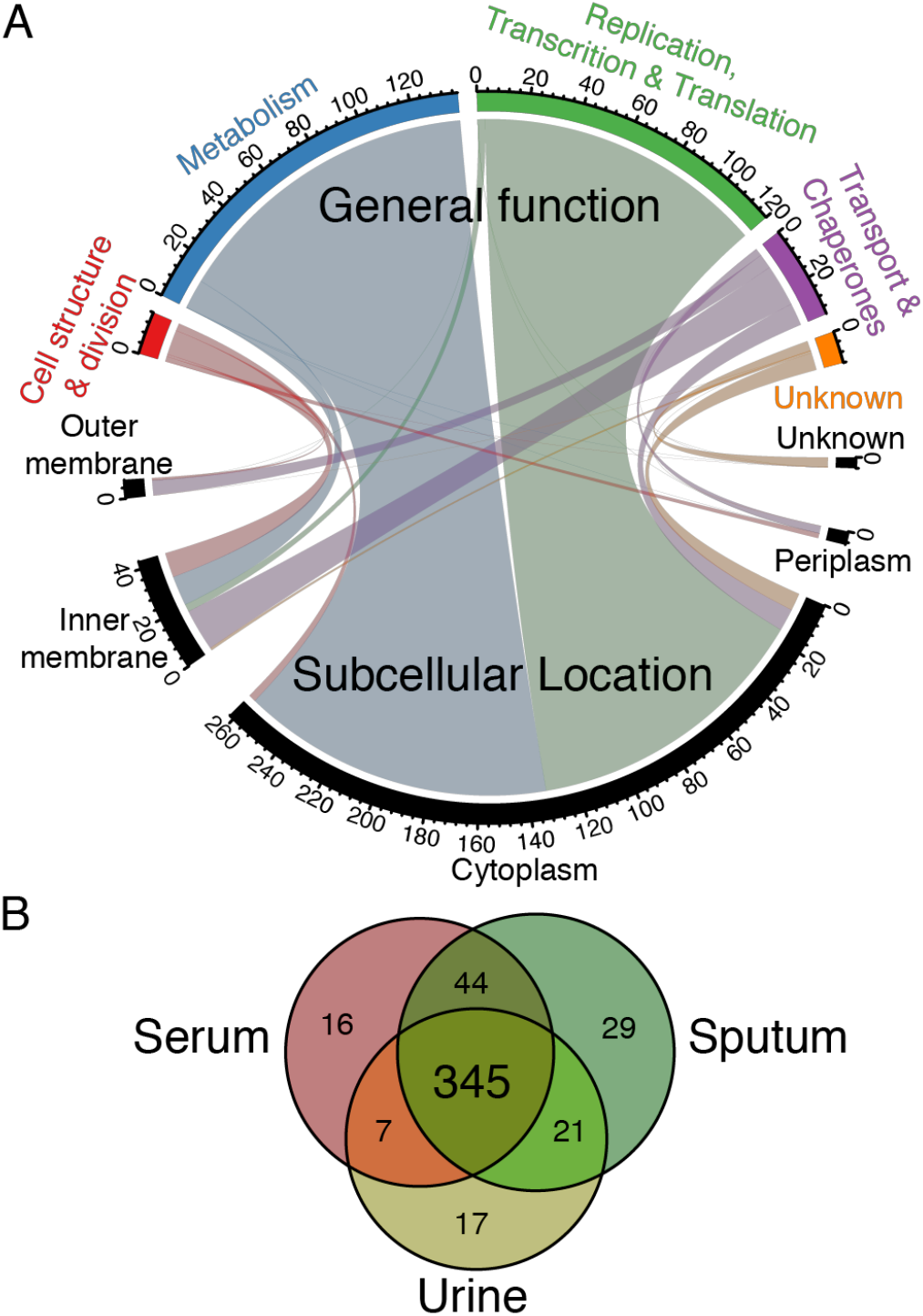
Core and conditionally essential gene functions in *P. aeruginosa*. A. Chord diagram of the 321 core essential genes showing the relationship between subcellular location (bottom) and general function (top), with the number of genes for each category indicated. B. Venn diagram showing the number of essential genes in all strains across three infection-relevant media.

### Conditionally essential genes

In addition to the core essential genes, the core genome also contains conditionally essential genes that are essential in one or more – but not all – media. Sputum and M9 had the highest number of conditionally essential genes (118 and 110, respectively), consistent with these being the most nutritionally depleted media. LB had 103 conditionally essential genes, while urine and serum had the fewest (69 and 91, respectively) (Table 1). While the numbers of essential genes required in each growth condition did not vary significantly from condition to condition, the actual gene identities did vary (Dataset S5). Importantly, we identified an additional 24 conditionally essential genes required for growth in all three infection-relevant media (serum, sputum, and urine; Fig. 4B) but not in both of the lab-based media (LB, M9). Several of these genes are involved in pyrimidine and purine synthesis and are not required in LB, suggesting that sufficient nucleotide intermediates may be present in LB to sustain growth *in vitro* but that these genes may be valid targets during *in vivo* infection.

When we applied Multiple Correspondence Analysis (MCA) to all sets of essential genes for every strain-growth condition, we find that indeed, the vast majority of strains formed distinct clusters based on growth condition (Fig. S6). Interestingly, one strain, PA14, an extensively used laboratory strain, was an outlier under two conditions, M9 and urine. This behavior could be a result of the strain simply having genetic idiosyncrasies that, for example, may contribute to its unique ability to colonize a wider host range than other *P. aeruginosa* strains (31, 32). Alternatively, this might be a consequence of PA14 being a laboratory strain which has adapted to laboratory conditions over a long period of time, perhaps providing a slight cautionary flag if attempting to extrapolate PA14 behavior to the species in general.

Contained within the sets of conditionally essential genes for each growth condition are genes that are essential only in a single medium, termed ***unique conditionally*** essential genes. Considering only the three infection-relevant conditions while ignoring the laboratory conditions, sputum had 29, serum had 16, and urine had 17 unique conditionally essential genes. These unique conditionally essential genes carry the intriguing potential of becoming infection site-specific targets for infection type specific antibiotics, for example, a urine specific anti-pseudomonal antibiotic, as long as their essentiality is not dependent on factors that are variable from patient to patient.

The essential genes unique to sputum consist mainly of biosynthetic pathways such as thiamine, pyridoxine, and tryptophan synthesis, with the former two cofactors being required for multiple cellular processes including synthesis and catabolism of sugars and amino acids and the latter requirement suggesting that tryptophan levels in sputum may not be sufficient for growth (23). Similarly, urine-specific essential genes almost exclusively consist of genes involved in amino acid biosynthesis, specifically valine, leucine, and isoleucine pathways. Meanwhile, methionine and arginine pathways are essential in both urine and serum. The urine findings are consistent with the fact that these amino acids are among the least abundant in urine (33). However, despite the low abundance of proline and cysteine in urine, classical proline and cysteine biosynthesis genes are not essential, likely because alternative, functionally redundant synthesis pathways exist in *P. aeruginosa* for these amino acids (34, 35).

In contrast, most genes involved in amino acid biosynthesis are dispensable in serum, as are genes involved in heme biosynthesis (*pdxA,H*, and *hemA,B,C,D,E,F,H,J,L*), likely due to the ability to scavenge amino acids (36) and heme from free hemoglobin (37, 38) from serum. Interestingly, despite the non-essentiality of porphyrin genes in serum, genes involved in the formation and utilization of porphyrin-containing cytochrome c were uniquely essential in serum and no other media, including the cytochrome c biogenesis protein CcmH (PA14_57540), Cytochrome c1 (Cyt1; PA14_57540), the ubiquinol cytochrome c reductase (PA14_57570), and cytochrome c oxidase cbb3-type subunit I (CcoN; PA14_44370). *P. aeruginosa’s* respiratory chain is highly branched and able to use diverse electron donors and acceptors under different environments (39). Here we find that in an environment containing high concentrations of heme, *P. aeruginosa’s* respiratory flexibility is lost, as it becomes dependent on a single pathway.

To validate *in vivo* a strategy of targeting conditionally essential genes identified from our *in vitro* Tn-Seq experiments, we tested a set of PA14 deletion mutants for their ability to survive in different *in vivo* mouse models of *P. aeruginosa* infection. Using a neutropenic mouse model where the bacteria are administered intravenously to test translation of the *in vitro* serum growth condition, we infected mice with 6 strains: wild-type PA14; 3 mutants containing deletions of metabolic genes predicted to be essential in serum, sputum, and urine but not LB (pyrC, pyrimidine biosynthesis; *tpiA*, glycolysis; and *purH*, purine biosynthesis); one mutant containing a deletion in a gene predicted be essential in serum and urine but not sputum (argG, arginine biosynthesis); and one mutant containing a deletion in a gene predicted to be conditionally essential in sputum alone but not serum or urine (thiC, thiamine biosynthesis; Fig. 5A). In concordance with their predicted conditional essentiality, the *pyrC, tpiA*, and *purH* mutants were significantly attenuated in the neutropenic mouse model with a 3-4 log reduction in total CFU in the spleen 16 hours post infection. Interestingly, the *argG* mutant was not attenuated as predicted. To understand this discrepancy, we compared its growth on agar plates supplemented with mouse, bovine (which was used in the Tn-Seq experiments), or human serum. Indeed, the mutant was able to grow on mouse serum, but not bovine or human serum, thereby explaining the lack of mutant attenuation in the mouse model, likely due to the higher levels of arginine available in mouse serum (40-42) (Fig. 5B). Finally, as predicted, the *thiC* mutant was not significantly attenuated in the neutropenic mouse model; it was however, attenuated in an acute pneumonia mouse model, where the inoculum is delivered intranasally followed by colonization of the lungs, which is consistent with *FiTnEss* predictions (Fig. 5C). Together, these datasets highlighted the tremendous differences required by *P. aeruginosa* in different microenvironments that could be exploited in the development of infection condition specific therapeutics.

**Figure 5.**
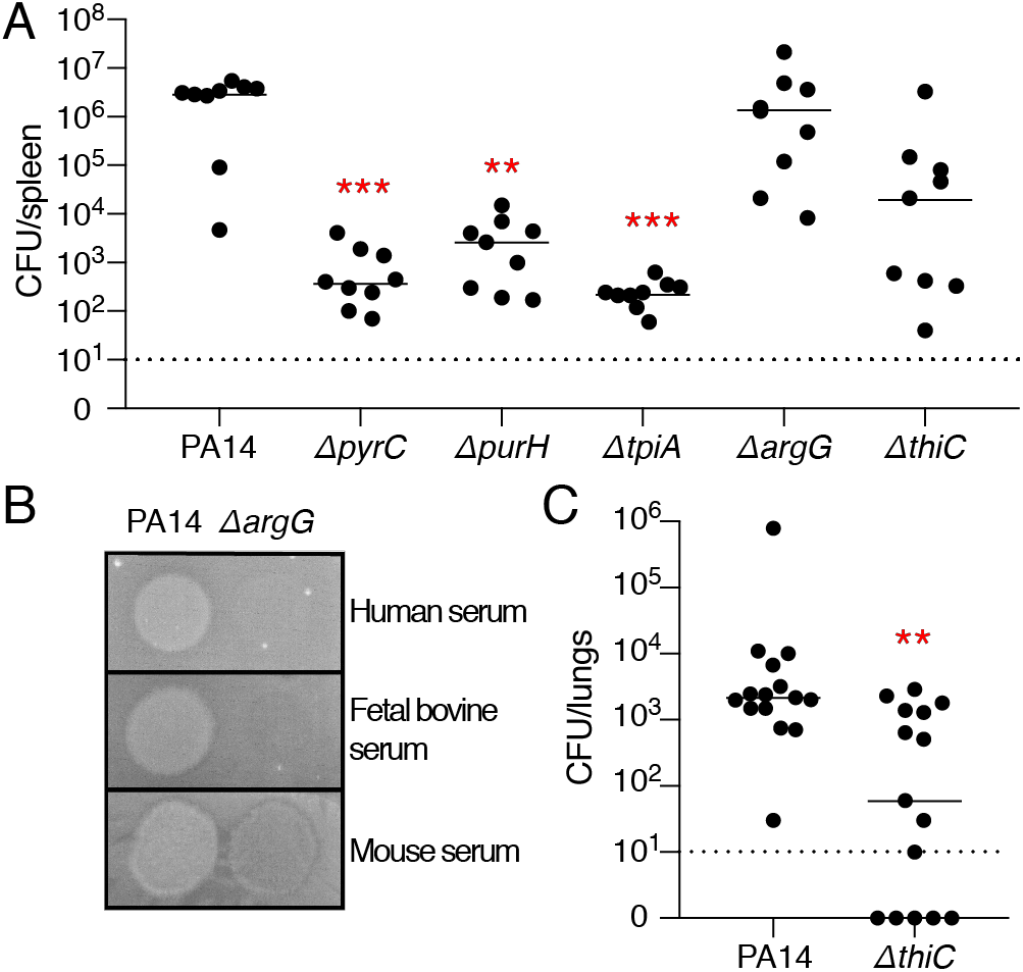
*In vivo* murine infection with gene deletion mutants of conditionally essential genes identified by *FiTnEss*. A. Bacterial burden in the spleen of neutropenic mice infected intravenously with WT PA14 and *ΔpyrC, ΔpurH, ΔtpiA, ΔargG*, and *ΔthiC* deletion mutants; statistical significance was determined with a Kruskal-Wallis test, n = 9. B. Growth of WT PA14 and the *ΔargG* deletion mutant in human, fetal bovine, and mouse sera. C. Bacterial burden in the lungs from mice infected intranasally with PA14 or the *ΔthiC* deletion mutant; statistical significance was determined with a Mann-Whitney test, n = 15. A and C. Bacterial burden determined at 16 hours post infection; each dot represents a single mouse with a line indicating the median; significance is displayed at p < 0.001 (***) or p < 0.01 (**); dashed line indicates the limit of detection of the assay; data is a combination of two biological replicates.

## Discussion

Target-informed antibiotic discovery and development has been predicated on knowing which genes within a given species constitute good targets, with genomic technologies such as Tn-Seq paving the way for more comprehensive definition of essential targets. However, comprehensive genomic methods for defining essential targets, such as Tn-Seq, have largely been applied to only a single or a few bacterial strains under a single or few growth conditions, with the implicit assumption that these results will apply to the entire species under infection-relevant conditions. Here we show that the number of essential genes for a given strain on a single media can vary widely among strains within a bacterial species and under different growth conditions. We thus sought to develop a robust paradigm for defining the core essential genes of a bacterial species, requiring an analysis of essential genes across multiple media and among multiple strains. We empirically determined that the analysis of essential genes among 4 strains was enough for *P. aeruginosa* to converge on a set of core essential genes that are likely to be essential in a wide range of conditions relevant to *in vivo* infection, and therefore represent the most attractive targets for novel drug discovery.

Using Tn-Seq and a novel method of analysis, *FiTnEss*, to establish the core essential genome of *P. aeruginosa*, we determined that while a single strain has ~400-800 essential genes, the core essential genome across all strains analyzed is approximately 321 genes, thus demonstrating the limitations in determining species essentiality based on a single strain. Further, there are an additional 24 essential genes required for growth in the three infection-relevant media examined, which are nonessential in LB and M9 media. Finally, we find that there are ~15-30 unique, conditionally essential genes for each of the infection-relevant media examined, with their corresponding biological pathways important for survival only within a particular host tissue environment; these genes may represent a unique set of targets for infection-type specific therapeutics with the obvious caveat that their essentiality cannot be dependent on microenvironmental factors that vary widely from patient to patient.

Previous transposon mutagenesis studies of two common lab strains, PA14 and PAO1, have found varying numbers of essential genes, as reviewed in (18). These studies have predominantly focused on identifying genes refractory to transposon mutagenesis when bacteria are selected for growth on lab media including LB, BHI, and minimal media (20, 43-46), though recent studies have extended growth conditions to include sputum or sputum-like media (45, 46) *in vitro*. A comparison of all of these datasets combined, revealed an intersection of only 109 essential genes common to all studies (Dataset S6). This low concordance is likely due to methodological or analytical differences between the studies. One way in which our study differs from the majority of previous studies is that we did not initially select transposon mutants on a rich (isolation) medium, *i.e.*, LB, prior to selection on the condition of interest, thereby eliminating a bottleneck that prevents evaluation of genes essential in the isolation media. By omitting the initial isolation step, we were able to identify 103 conditionally essential genes which are required in LB, but not in at least one of the other 4 media. In addition to these *in vitro* studies, *in vivo* Tn-Seq studies can be valuable in determining what genes are required for fitness in the context of an active infection (44); however, they also suffer from the problem of experimental bottlenecks that practically limit the ability to truly interrogate essentiality on a genome-wide scale. These bottlenecks include the required isolation step before inoculating into an animal and the depletion of mutants *in vivo* due to stochastic loss rather than a true fitness loss of the mutant itself (13).

Experimental methods for identifying gene essentiality have varied greatly through the years. Despite significant advances for defining fitness costs of gene disruptions on a genome-wide scale using sequencing (Tn-Seq), limitations persist. They include (1) analysis of mutant behavior in pools where there can be both competition as well *trans* complementation, factors which can cause mutant growth in a pool to diverge from growth in isolation, (2) transposon sequence insertion biases (24, 47), and (3) polar effects on adjacent genes conferred by transposon insertion. These limitations prevented us from evaluating the essentiality of approximately 5% of the genome and can technically lead to errors in assessing fitness and essentiality. We indeed found examples of discordance where our Tn-Seq data classified genes as essential even though mutants of these genes are available, albeit growth defective, such as *hfq, rpoN*, and *gidA* (48-50). These cases could be due to *FiTnEss* errors; alternatively, one must consider the possibility that these mutants have reduced fitness when grown in competition but not in isolation, or that reported mutants could contain compensatory mutants acquired in their construction that allow deletion of the gene of interest.

A major challenge to analysis is translating measurements that quantify a continuum of fitness to a binary classification of essentiality versus non-essentiality, in order to define the best antibiotic targets. Approaches can vary substantially, with different systematic errors and different tolerances for false positive versus false negative predictions (Fig. S7 and SI methods). We therefore developed a Tn-seq analysis pipeline, *FiTnEss*, that balances false positive and false negative rates with the aim of accurately classifying gene essentiality, while providing two levels of stringency depending on one’s tolerance for false positive versus false negative predictions. We validated *FiTnEss* using clean gene deletions mutants (Fig. 2 and Fig. S4).

We used *FiTnEss* to perform the binary classification of the 4903 genes in the core genome that could be assessed across 9 *P. aeruginosa* isolates. The great majority of core essential genes can be broadly categorized as being involved in metabolic pathways or macromolecular synthesis such as DNA replication, transcription or translation. That the core essential genome is dominated by genes involved in macromolecular synthesis (*i.e*. protein and nucleic acid) may explain in part why most current antibiotics seem to target this limited set of functions. There has been greater reticence to target metabolic pathways given concern over the ability of bacteria to scavenge nutrients from the host, thereby rendering their biosynthesis nonessential during infection. Indeed, we see variable requirements for metabolic genes in our identification of conditionally essential genes, such as a greater dependence on amino acid biosynthesis in urine than other growth conditions. We have however, demonstrated that this conditional essentiality can in fact be exploited *in vivo*, as mice infected with the *pyrC, tpiA*, and *purH* mutants that are significantly growth impaired on infection-relevant media, have dramatically reduced bacterial burden in a systemic infection model. Further, the *thiC* mutant, found to be essential in M9 and sputum alone, was attenuated in an acute pneumonia model, yet was still virulent if introduced systemically, demonstrating variable metabolite levels in different infection sites *in vivo* and raising the possibility of infection-site specific therapeutics.

In summary, we suggest the apparent failure of genomics to transform antibiotic discovery in the late 1990s to early 2000s was due not to a fundamental flaw with the concept of targeting essential genes, but rather with challenges in implementing the approach — namely, defining essential genes based on limited information. Advances in genomic technologies now make possible studies on a much greater scale, allowing us to define essential genes in a way that overcomes previous shortcomings. Our work describes a general approach applicable to other pathogens, given the explosion in available bacterial genomes. While the number of strains required to reach a plateau in essential genes for different species may vary based on the genomic diversity of a species, the basic paradigm should apply broadly. By selecting diverse strains across the phylogenetic tree of any species, renewed efforts to identify the essential genes in all major bacterial pathogens may allow us to more effectively work towards the discovery and development of new, much-needed antibacterial therapeutics.

## Materials and Methods

### Strain selection and plasmid construction

A genome tree report of 2560 sequenced *P. aerguinosa* strains was downloaded from NCBI (organism ID: 187) and visualized with iTOL (51). Nine strains were selected for genetic diversity and graciously gifted from various sources: PA14, 19660, X13273 obtained from Frederick M. Ausubel (52); BWH005, BWH013, BWH015 were collected through Brigham and Women’s Hospital Specimen Bank per protocol previously described (53); BL23 from Bausch & Lomb (54); PS75 from Paula Suarez, Simon Bolivar University, Venezuela; and CF77 from Boston Children’s Hospital (55). pC9 containing a hyperactive transposase was derived from pSAM-DGm (44) and pMAR containing the Himar1 transposon was derived from pMAR2xT7 (20).

### Transposon library construction and sequencing

Recipient *P. aeruginosa* strains were prepared for mating as previously described (56). *P. aeruginosa* and mid-log cultures of *E. coli* SM10(pC9) and *E. coli* SM10(pMAR) were collected by centrifugation, washed and resuspended in LB. A total of 3 × 10^11^ CFU were mixed in a 2:2:1 ratio of pC9:pMAR:recipient and collected by centrifugation. The cell mating mixture was re-suspended to a concentration of 10^10^ CFU/ml and 30 μl spots were dispensed to a dry LB agar plate. Mating plates were incubated at 37°C for 1.5 hours before cells were scraped, resuspended in phosphate buffered saline (PBS), mixed with glycerol to a final concentration of 40%, aliquoted, and flash frozen before storage at −80°C. Matings were performed at least twice for each recipient strain and efficiencies were quantified by plating to LB selective agar. 250 mL of each medium containing 1.5% agar, 5 μg/ml irgasan, and 30 μg/ml gentamicin was prepared in a Biodish XL (Nunc). LB and M9 minimal agar (US Biologicals), and synthetic cystic fibrosis medium agar (SCFM) (23) were prepared. Pooled, filter-sterilized urine, and fetal bovine serum (FBS) (ThermoFisher) were warmed to 55°C and mixed with a 5% agar solution (Teknova) to achieve a 1.5% final agar concentration. 500,000 CFU of each transposon-integrated strain were plated to each medium in duplicate and grown at 37°C for 24 hours (LB, FBS, SCFM) or 48 hours (urine, M9) before scraping and re-suspending cells in PBS. Genomic DNA was isolated and Illumina libraries were prepared using a custom method described in the SI (Fig. S1 and Dataset S7). Sequencing was performed with an Illumina Nextseq platform to obtain 50 bp genomic DNA reads.

### Determining essential genes from Tn-Seq data using *FiTnEss*

All software scripts are freely available (https://github.com/ruy204/FiTnEss). Genomes and annotations for each strain were obtained from www.pseudomonas.com and www.patricbrc.org (57). The core and accessory genomes were determined by gene clustering analysis across the strains tested using Synerclust (29). Illumina reads were mapped to each respective genome using Bowtie (58) using the options for exact and unique read mapping. Reads potentially mapping to more than one location in a genome were discarded and homologous TA sites were removed from analysis by searching the genome using custom scripts. TA insertion sites at the distal 50bp from each end of the gene and non-permissive insertion sites containing the sequence (GC)GNTANC(GC) were removed using custom scripts. Reads mapped to each TA site were tallied using scripts modified from (28). For each Tn-Seq dataset, a lognormal – negative binomial distribution was conservatively fit using genes with median number of TA sites, and top 75% of reads per gene (non-essential genes) to identify parameters (*μ*, *σ*). Then a theoretical distribution 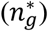 was constructed using these two parameters for each gene size category based on the number of TA sites per gene (*N_TA_*). Background distributions for these categories were obtained from numerical sampling of the theoretical distribution. The actual number of reads for each gene was compared to the background distribution for the corresponding *N_TA_* category, and a p-value was calculated as the probability of obtaining the number reads (*n_g_*) or less by chance. Two-layer multiple comparison adjustments were conducted. First, to obtain a maximally stringent essential set, we adjusted for family-wise error rate (FWER) using the Holm-Bonferroni method. Second, to reduce the risk of losing targets we relaxed the stringency slightly to obtain a highly stringent essential set, by adjusting for false-discovery rate (FDR) using the Benjamini-Hochberg method. After either correction process, genes with adjusted p-value smaller than 0.05 in both replicates are identified as essential. For a full description and calculations please see the SI.

### Method validation with clean gene deletions

Gene deletions were performed as previously described in strain PA14 (56). Gene deletions were confirmed by PCR amplification and sequencing. Successful gene deletion strains were grown in duplicate in LB at 37°C for 16 hours before diluting 10^−4^ in PBS. 5 μl diluted culture was spotted to the five solid media used in this study and grown at 37°C for 24 hours. Images were captured, densitometry was performed using ImageJ, and growth was categorized relative to 10 wild type replicates: essential (0-20%), growth-defective (21-50%), and nonessential (>50%).

### *In vivo* mouse models

All vertebrate animal experiments were done with with the approval of Massachusetts General Hospital’s Institutional Animal Care and Use Committee. Bacteria were grown to mid-log, collected by centrifugation, washed and resuspended in PBS. For the systemic infection model, 9 week old female BALB/c mice (Jackson Laboratory) were injected intra-peritoneally with 4 mg cyclophosphamide 3 days prior to infection. Mice were infected intravenously with 5 × 10^5^ CFU per mouse. For the acute pneumonia model, mice were infected intranasally with 1 × 10^6^ CFU per mouse. For both infection models, mice were euthanized 16 h post infection and spleens (systemic) or lungs (pneumonia) were harvested and homogenized in 1 mL PBS + 0.1% Triton-X100 using a TissueLyser LT (Qiagen) before plating to LB agar + 5 μg/ml irgasan to enumerate bacterial burden.

## Acknowledgements

This work was supported by a generous gift from Anita and Josh Bekenstein, NIH grant R33AI098705 (DTH), and a Cystic Fibrosis Canada Fellowship (BEP).

