## Supplementary Information for "Defining the core essential genome of *Pseudomonas aeruginosa*"

### Supplementary information (datasets, methods and results, figures, tables)

**Datasets**

Dataset S1 - sequencing depth and replicate concordance.xlsx
Dataset S2 - genes without usable TA sites.xlsx
Dataset S3 - gene probabilities of essentiality for all strains and media.xlsx
Dataset S4 - 13 potential core essential genes.xlsx
Dataset S5 - core and conditionally essential genes in core genome.xlsx
Dataset S6 - core essential genes comparison with previous studies.xlsx
Dataset S7 - primers used in this study.xlsx

Datasets (.xlsx Microsoft Excel files) can be found here:
https://drive.google.com/drive/folders/1EFF3y-I1UGNP3dRa5PMmSL7qcXrNqqav?usp=sharing

### Additional Methods and Results

**Plasmid construction.** pC9 was derived from pSAM-DGm (1) by digesting with ApaLI + AccI to remove the transposon, and pMAR was derived from pMAR2xT7 (2) by digesting with ApaLI + StuI to remove the transposase. Linearized vectors were each blunted, phosphorylated, ligated, and transformed into *E. coli* SM10 donor cells and selected on 100ug/ml carbenicillin (pC9) or 15ug/ml gentamicin (pMAR). Cloning reagents were obtained from New England Biolabs.

**Transposon library construction and sequencing.** Overnight cultures of *E. coli* SM10(pC9) and *E. coli* SM10(pMAR) donor cells were grown in LB medium with their respective antibiotics, sub-cultured 1:100, and grown at 37°C while shaking at 250 RPM for 3.5 hours until OD600nm reached ~0.5. Overnight cultures of recipient *P. aeruginosa* strains were grown in LB medium, sub-cultured 1:5, and grown at 42°C while shaking at 250 RPM for 3.5 hours. Cells were collected by centrifugation at 5000g for 10 minutes, washed once, and re-suspended in LB. A total of 3 x $10^{11}$ CFU were mixed in a 2:2:1 ratio of pC9:pMAR:recipient and collected by centrifugation. The cell mating mixture was re-suspended to an approximate concentration of $10^{10}$ CFU/ml and 30 µl spots were dispensed to a dry LB agar plate. Mating plates were incubated at 37°C for 1.5 hours before cells were scraped, resuspended in phosphate buffered saline (ThermoFisher), mixed with glycerol to a final concentration of 40%, aliquoted, and flash frozen in a dry ice/ethanol bath before storage at -80°C. A small aliquot of each mixture was thawed, diluted and plated to 5 µg/ml irgasan + 30 µg/ml gentamicin for CFU quantification of successful integrants. Matings were performed at least twice for each recipient strain. 250 mL of each medium containing 1.5% agar, 5 µg/ml irgasan, and 30 µg/ml gentamicin was prepared in a Biodish XL (Nunc). LB agar (US Biologicals), M9 minimal agar, synthetic cystic fibrosis medium (SCFM) (3) were prepared as previously described. Pooled, filter-sterilized urine, and fetal bovine serum (FBS) (ThermoFisher) were warmed to 55°C and mixed with a 5% agar solution (Teknova) to achieve a 1.5% final agar concentration. 500,000 CFU of each transposon-integrated strain were plated to each medium in duplicate and grown at 37°C for 24 hours (LB, FBS, SCFM) or 48 hours (urine, M9) before scraping and re-suspending cells in PBS. Genomic DNA was isolated using the DNeasy kit (Qiagen), and 5 µg from each sample was sheared to 1.5 kb fragments by sonication (Covaris). End repair, dA-tailing, P5 adapter ligation, and PCR of the transposon-gDNA junction was performed using NEBNext enzymes (NEB) and custom primers from IDT (Fig. S1 and Dataset S7). Size selection was performed using Agencourt Ampure XP beads (Beckman Coulter) and ~500 bp libraries were quantified using D5000 ScreenTape System (Agilent). Sequencing was performed with an Illumina Nextseq platform to obtain 50 bp genomic DNA reads.

**Removal of confounding transposon insertion sites from analysis.** At these TA sites, the presence or absence of mapped insertions can be influenced by methodological artifacts unrelated to the essentiality of gene in which the TA is located. To avoid these confounding factors, we removed three classes of TA sites from analysis because of their potential to mislead:

(1) Non-permissive insertion sites – The sequence (GC)GNTANC(GC) was reported to be intolerant to Himar1 transposon insertions in *Mycobacterium tuberculosis* (4), which has a similar GC content to *P. aeruginosa*. This sequence occurs 6367 times in *P. aeruginosa* strain PA14 across 3389 genes. Indeed, we found that insertions mapped to these sites at a significantly reduced frequency compared to a random subsample of TA sites ($p < 0.0001$, Fig. S2) and thus excluded them from all subsequent analysis.

(2) Non-disruptive terminal insertions – Transposon insertions close to 5' and 3' gene termini can nevertheless result in the expression of a functional, albeit truncated version of the corresponding gene product (5). Rather than selecting an arbitrary distance from the termini in which to exclude such potentially confounding TA sites, we empirically determined an optimal distance. Using the consensus

109 essential genes from previous transposon studies of strains PA14 and PAO1 as the truth set for essential genes (1, 2, 6-8), we found that 42 of these genes in our PA14-LB dataset contained >10 sequencing reads, all of which corresponded to TA site insertions located within 50 bp from the gene termini, regardless of gene size (Fig. S2). We thus eliminated from analysis all TA sites that fell within 50 nucleotides of either the 5' or 3' ends of each gene. Removal of these confounding TA sites resulted in the exclusion of 9829 TA sites.

(3) Homologous insertion sequences – Because insertions are assigned to a specific TA site in a specific gene based on the mapping of the genomic sequences flanking the ends of a transposon onto the entire genome, we removed from consideration TA sites whose flanking regions are not unique because of the possibility of mis-mapping reads.

***FiTnEss*: a statistical model to identify essential genes.** After removal of confounding TA sites, we calculated the total number of reads **ng** by summing over the reads at all remaining TA sites in the gene, where **NTA** is the number of TA sites in the gene. By calculating average number of reads per TA site for a gene (**ng/NTA**), we observed a clear bimodal distribution for genes containing more than a few (≥5 TA) sites (see NTA = 5,10,15 in Fig. S3AB). Presumably, in these distributions, essential genes are on the left with a small or zero ng/NTA, and non-essential genes are on the right (ng/NTA>0). We noted that ng/NTA was not the same for all NTA categories, with the standard deviation decreasing (unsurprisingly) as the number of TA sites increased (Fig. S3E), while the distribution of reads at single TA sites was similar for all NTA categories (Fig. S3CDF). This suggested that all TA sites were behaving the same, independent of gene length. Given these bimodal distributions, we based *FiTnEss* on modelling of the read-number distribution for non-essential genes and fitting the model parameters from the data (see below).

We posited that the distribution of the number of reads at any TA site in a gene is geometric with probability pg, and that the expected number of reads (1/pg) is only a function of the fitness of the bacteria when the gene function is lost. Assuming a lognormal distribution of 1/pg, the model only requires two parameters, the mean and variance of this distribution, to be determined from the data. Since the model describes non-essential genes, it was important to avoid data from essential genes when fitting the model parameters. Thus, we used only genes with which we had high confidence in their non-essentiality (NTA = 10; top 75% of the distribution). Given the fitted parameters of the model, a specific dependence of the distribution of ng on the number of TA sites was predicted, which agrees well with the actual data (red curves in Fig. S3B).

**A model for non-essential genes.** We assume that each non-essential gene g is characterized by a parameter, $p_g$, the inverse of which comes from a log-normal distribution

$$p_g{}^{-1} \sim Lognormal(\mu, \sigma), \qquad (1)$$

with parameters $\mu, \sigma$. We further assume that for any non-essential gene g with certain number of TA sites ($N_{TA}$), the read counts at any of its TA sites, $x_{g,i}$, are iid, and are distributed according to

$$\text{For a specific gene g: } x_{g,i} \sim Geo(p_g), \quad for\ i = 1, \ldots, N_{TA}. \qquad (2)$$

A possible interpretation of this model is that there is a distribution among non-essential genes of the small fitness costs of disabling them. Genes that are slightly more important would have a higher knockout cost $p_g{}^{-1}$, or a lower $p_g$, and thus a lower number of reads per TA site on average.

It follows that the distribution of $n_g$, the total number of reads in a given gene, follows a negative binomial distribution:

For a specific gene g:

$$n_g \equiv \sum_1^{N_{TA}} x_{g,i} \sim NB(N_{TA}, p_g). \tag{3}$$

The distribution of $n_g$ among all the genes for some value of NTA is the convolution of the lognormal and the negative binomial:

$$F_{n_g}^*(n) \equiv Prob(n_g \leq n) = \int_0^{+\infty} f_{LN}(\tfrac{1}{p}; \mu, \sigma) F_{NB}(n; N_{TA}, p) d\left(\tfrac{1}{p}\right), \tag{4}$$

where $f_{LN}$ is the probability density function of the lognormal distribution and $F_{NB}$ the negative binomial cumulative distribution function.

**Fitting model parameters.** Cramér-von Mises criterion is a goodness of fit criterion, measuring the difference between cumulative density functions of an empirical distribution and a fitted, theoretical one. We use it here for the distributions of the total number of reads in a gene with $N_{TA}$ TA sites (Equation 4). The empirical distribution $F_{N_{TA}}$ is obtained directly from the data, and we use numerical sampling to approximate the theoretical $F_{N_{TA}}^*$ (sampling 100,000 times for each pair of parameter values $\mu, \sigma$). For any $N_{TA}$ category, we have

$$\omega_{N_{TA}}^2 = \int_0^{+\infty} \left[F_{n_g}(n; N_{TA}) - F_{n_g}^*(n; N_{TA})\right]^2 dF_{n_g}^*(n; N_{TA}), \tag{5}$$

with $\omega_{N_{TA}}^2$ denoting the integral of squared distance between two functions for all genes with $N_{TA}$ TA sites. In order to fit model parameters which describe non-essential genes, we tried to avoid data that are potentially "contaminated" by essential ones in the parameter estimation phase. To address this, we use a modified version of the Cramér–von Mises criterion $\omega^2$ as follows:

$$\omega_{N_{TA}}^2 = \int_{n_{[1/4]}}^{+\infty} \left[F_{n_g}(n; N_{TA}) - F_{n_g}^*(n; N_{TA})\right]^2 dF_{n_g}^*(n; N_{TA}), \tag{6}$$

where $n_{[1/4]}$ is the low 25 percentile of read counts in this $N_{TA}$ category. This practically means that minimizing this distance is only affected by the goodness of fit to the higher 75% of the empirical distribution, which is not expected to contain essential genes.

The model parameters can be determined by minimizing the sum of this modified $\omega^2$ for any of the different $N_{TA}$ categories, and the resulting parameters are not affected much by this choice. Yet we observed that for genes with a low number of TA sites there was not much separation between the essential and non-essential populations. Conversely, for the gene categories with large $N_{TA}$, where this separation is very pronounced, the number of genes in these categories is too small and leads to less robust fits. We have estimated the variability of the fitted parameters under perturbations of the data, and concluded that using values of $N_{TA}$ between 5 and 15 yield robust fits (Fig. S8). The parameters used for the results in this paper are based on fitting the distributions at $N_{TA} = 10$.

**Calling essential genes.** For each Tn-Seq dataset (= a replicate of strain x medium), after identifying parameters $\mu, \sigma$ for non-essential genes, we construct the background distribution for each $N_{TA}$ category by sampling $10^5$ observations of $(n_g^*)$ from the theoretical distribution (Equation 4). The actual number of reads for each gene is then compared to the background distribution for the corresponding $N_{TA}$ category, and a p-value is calculated as the probability of obtaining this number reads ($n_g$) or less "by chance":

$$p\text{-}value = P(n_g^* < n_g). \qquad (7)$$

In each medium and strain, we have more than 5000 genes being tested simultaneously. Accounting for multiple testing is required for obtaining true signals. Two-layer multiple comparison adjustments were conducted. First, in order to obtain a more conservative essential set, we adjusted for family-wise error rate. Family-wise error rate (FWER) is a conservative correction method for multiple hypothesis, by controlling type I error we allow low probability of making one or more false discoveries. In our analysis, we used Holm-Bonferroni method with type I error rate $\alpha = 0.05$, indicating that we have only 5% chance of obtaining even a single false positive call in the dataset. Second, to reduce the risk of losing important targets by being too conservative, we used Benjamini-Hochberg procedure, which is a less strict approach controlling for false-discovery rate. After either correction process, genes with adjusted p-value smaller than 0.05 in both replicates are identified as "confident essential" (FWER) or "candidate essential" (FDR).

**Gene deletions.** Gene deletions were performed as previously described in strain PA14 (9). Briefly, 800-1200 bp regions flanking the target deletion gene of interest were PCR amplified, stitched and recombined into the pEXG2 (9) plasmid containing GentR and SacB markers using Gateway Cloning. Plasmids were conjugated into PA14 using the *E. coli* helper plasmid pRK2013 for 8 hours, followed by selection on LB agar containing 15 µg/ml irgasan + 30 µg/ml gentamicin. Individual colonies were grown in liquid LB for 4 hours, followed by streaking to LB agar supplemented with 10% sucrose and grown at 37°C for 16 hours. Colonies were confirmed to be GentS and successful gene deletions were confirmed by PCR amplification and sequencing.

**Comparison of *FiTnEss* to a HMM.** Analytical tools can also vary significantly, as they all have different strengths and weaknesses, often having been developed to answer different questions. One of the greatest challenges for the analytical tools is to translate measurements from Tn-Seq, which is really quantifying a continuum of fitness – from optimal growth in a particular condition to slow growth, from static for growth to cell death – to a binary classification of essential versus non-essential in the service of comprehensively defining candidate targets for antibiotic discovery. The different tools can vary dramatically both in the assumptions built into the analysis and how conservatively each model calls essentiality *i.e.,* whether one is more willing to tolerate false positives or false negatives. For example, a Hidden Markov Model (HMM) and sliding window approach rely on a stretch of TA sites that have zero to very low level insertions to denote an essential gene (10, 11). An advantage of these methods is that intergenic regions and essential domains within a larger gene can be queried. The disadvantage is that genes containing more insertions than the HMM and sliding window approaches tolerate in an essential gene, can in fact be essential with detectable insertions resulting because death of the corresponding mutant is slow or delayed. Because *FiTnEss* considers genes rather than individual TA sites as the basic unit for determining essentiality, it leverages all TA sites in a gene, allowing it to more easily distinguish whether low insertion numbers are indicative of low coverage in a non-essential gene or background noise in an essential gene. Another set of genes that are often discrepant between analytical methods is short genes that may be flanked by genes of the opposite classification. Here, approaches that examine "windows" of adjacent TA sites (~5-10 adjacent sites) can misclassify the short gene of interest (<5 TA sites) by erroneously integrating in data from the flanking genes which are of the opposite classification; because *FiTnEss* examines the gene independent of flanking regions, it can avoid being misled by the behavior of the TA sites in the flanking genes. In the particular case of longer genes containing a mix of essential and nonessential domains, the power of *FiTnEss* to detect essentiality is reduced because of it cannot distinguish the even distribution of reads across the gene (resulting in a call of non-essentiality) with a bimodal distribution of reads among the essential and non-essential domains (which should result in a call of essentiality). Here, other methods such as HMM outperform *FiTnEss* (Fig. S7 for comparison of methods, Dataset S3 for complete *FiTnEss*/HMM gene calls). While the methods are complementary, in the analysis of the datasets generated in this study, *FiTnEss* seemed to be generally more powerful for calling essential genes than HMM, with greater accuracy in calling the 115 conditionally essential gene-growth condition combinations that we validated using clean genetic deletions (Fig. 2 and Fig. S4). While the HMM method did not have any false positives (if combining essential and growth defective categories), it did miss calling many essential genes, *i.e.* tolerated a high false negative rate (Table S2). Meanwhile, *FiTnEss* attempted to balance false positive and false negative rates for this limited set of deletions resulting in greater overall accuracy. Of note, the genes selected for validation were skewed towards falling relatively clearly in the essential or non-essential distributions; thus they may overestimate the positive predictive power of *FiTnEss*, particularly for genes that lay at the boundary of the bimodal distribution. Nevertheless, overall, *FiTnEss* appears to perform well in its binary classification of genes.

### Figures

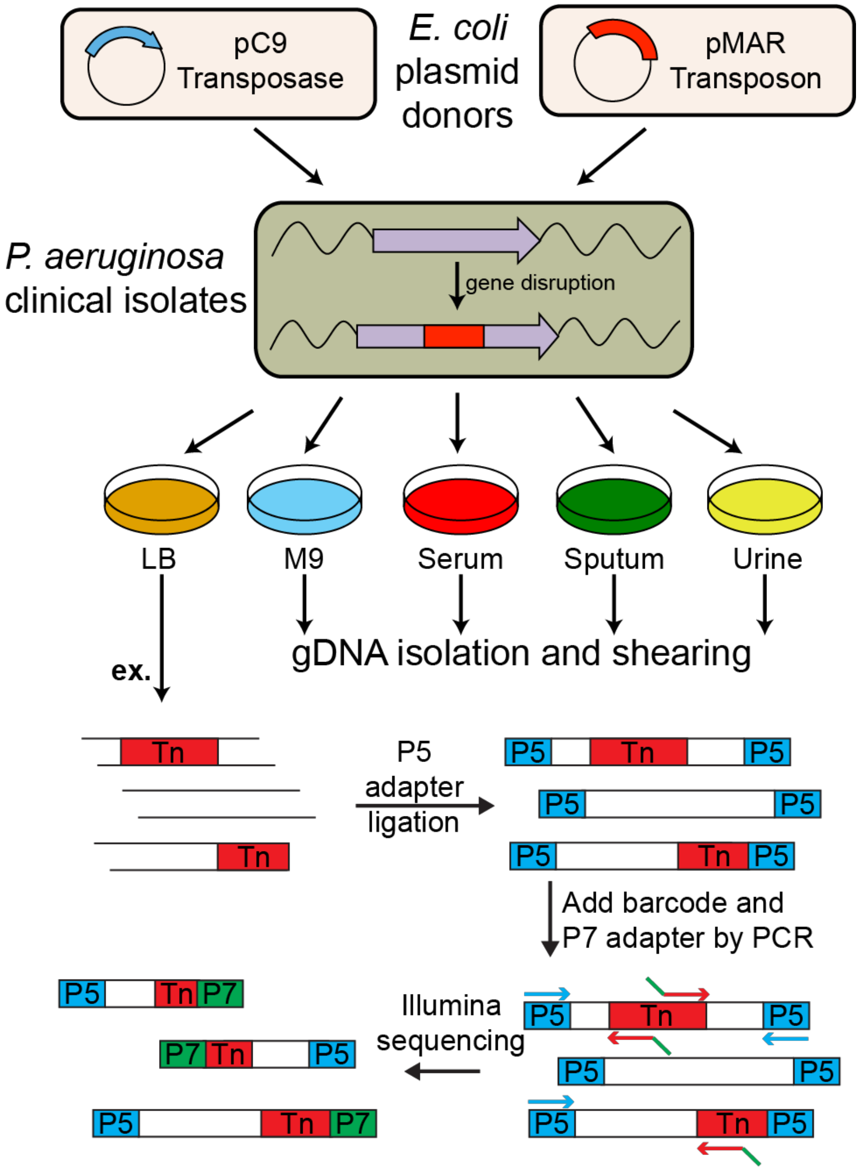


**Figure S1. Transposon mutagenesis and Illumina library construction.** *E. coli* SM10 donor cells containing either the pC9 transposase or pMAR transposon are mated with recipient *P. aeruginosa.* Transposon-integrated *P. aeruginosa* mutants are selected on solid medium: LB, M9 minimal, fetal bovine serum, synthetic cystic fibrosis sputum or urine followed by outgrowth, cell collection, and genomic DNA purification. Illumina libraries of isolated genomic DNA were constructed by: 1) end repair of ~1.5kb sheared DNA and ligation of Illumina P5 adapters; 2) PCR amplification using primers specific to the P5 ligated ends and transposon (Tn) while introducing the P7 Illumina flow cell binding site sequence; 3) size-selection of 400-500bp fragments containing the genomic-transposon junction.

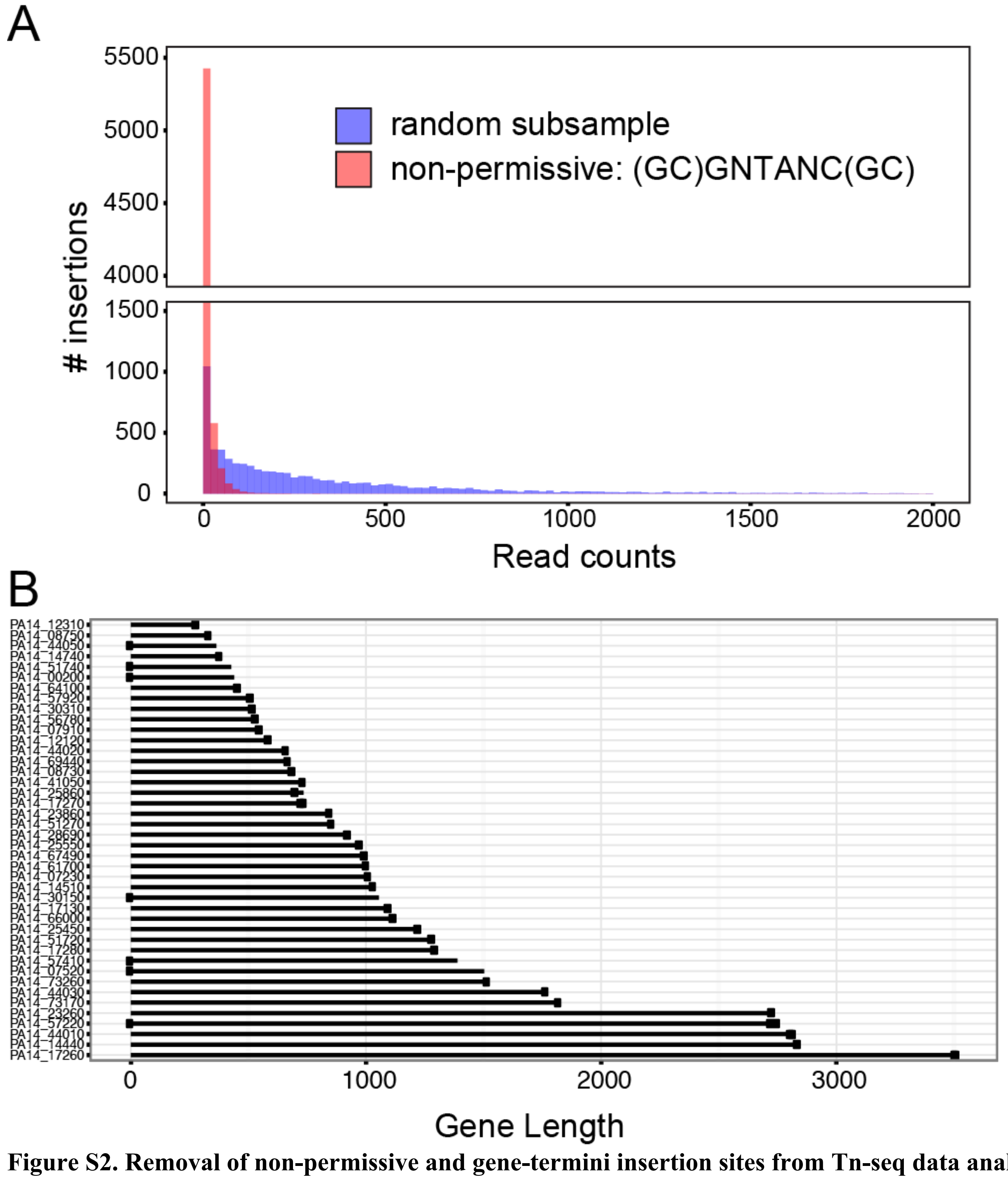


**Figure S2. Removal of non-permissive and gene-termini insertion sites from Tn-seq data analysis.** A. Non-permissive TA sites. Reads at TA sites with the surrounding sequence (GC)GNTANC(GC) have significantly reduced reads compared to a random subsample of TA sites ($p < 0.0001$, $n = 6352$, Kolmogorov-Smirnov test); PA14 LB dataset. B. Reads at termini of essential genes. 42 genes identified from our Tn-seq data that contain reads. These 42 genes are part of the consensus of 109 published essential genes (Dataset S6) and contain >10 reads from the PA14-LB dataset, as indicated with a dot. Reads in these essential genes are found <50 bp from either the 5' or 3' end regardless of gene size, thus distal TA sites were removed from analysis in all genes.

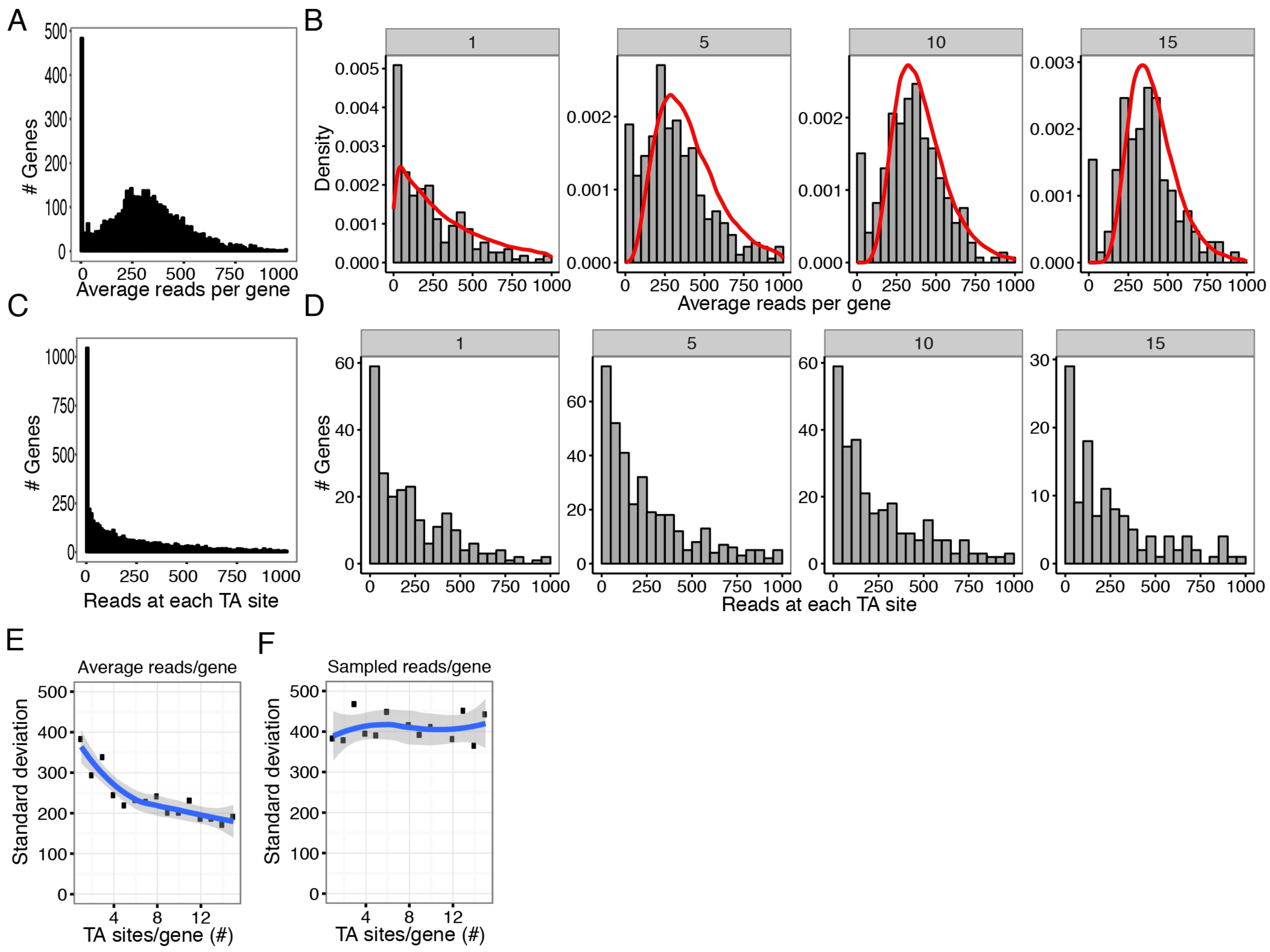


**Figure S3. Distributions of read numbers in Tn-Seq data in relation to gene size.** A. Distribution of average number of reads/TA site in a gene (ng/NTA) for all genes and B. in genes with only 1, 5, 10, 15 TA sites. The red curves are theoretical distributions for the non-essential genes simulated from our parameters; actual Tn-Seq data shown as histograms. C. Distribution of number of reads at one random sampled TA site in a gene for all genes and D. in genes with only 1, 5, 10, 15 TA sites. E. Standard deviation of average number of reads/TA site in a gene for these NTA categories is decreasing, as expected with increasing numbers of TA sites. F. Standard deviation of number of reads at one random TA site is relatively constant across different numbers of TA sites, thus showing that all TA sites are behaving similarly, regardless of gene length and numbers of TA sites in a gene.

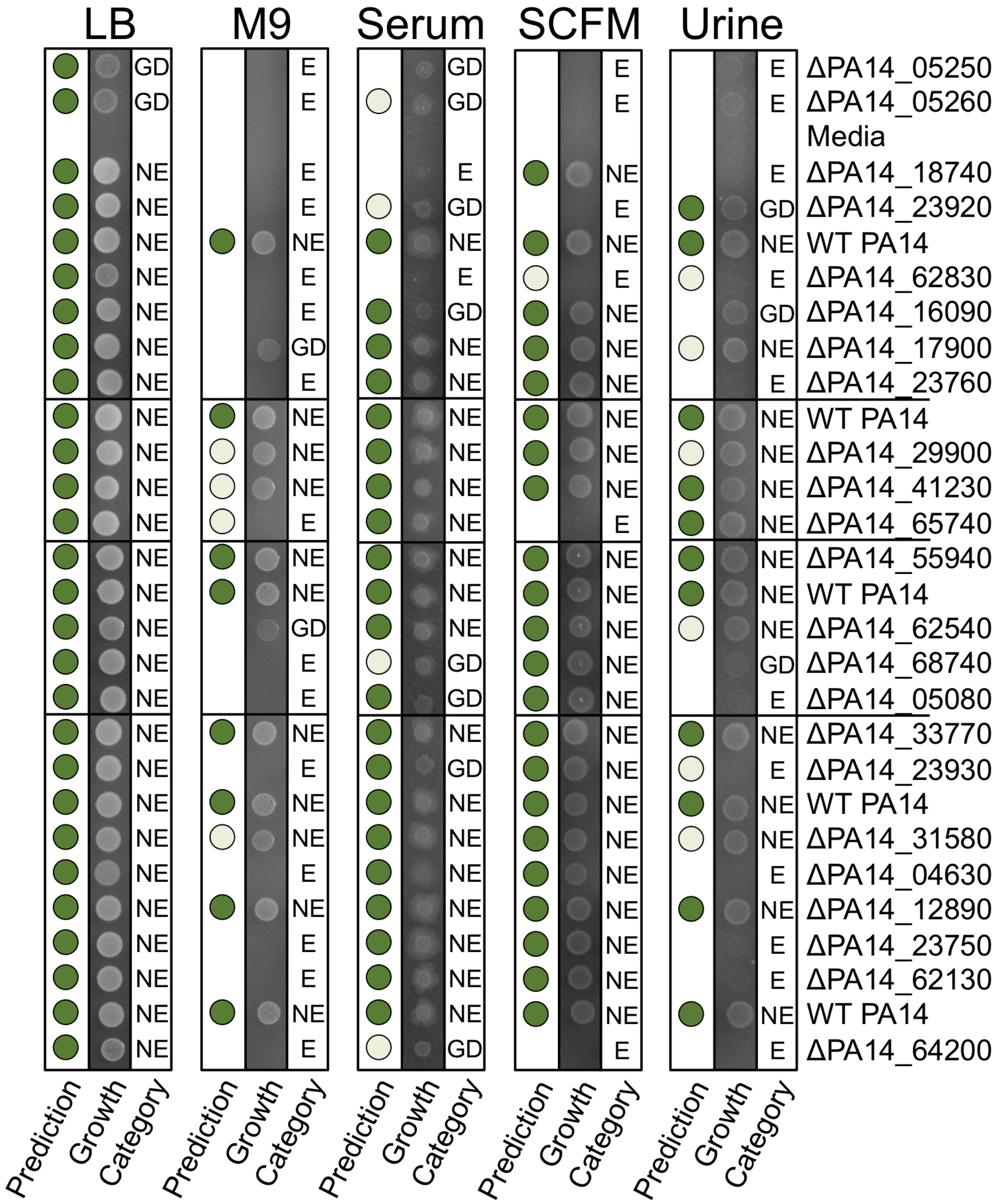


**Figure S4. *FiTnEss* validation using conditionally essential gene deletions.** 23 gene deletions in strain PA14 grown on 5 media (left to right: LB, M9, serum, CF sputum, urine). *FiTnEss* essentiality predictions (left columns) are depicted by: dark green circle, non-essential; no circle, confidently essential; light green circle, candidate essential. Images of growth (middle columns) are categorized using densitometry of 2 biological replicates relative to 10 replicates of WT (right columns): E = 0-20% of WT, GD = 21-50% of WT, NE > 50% of WT.

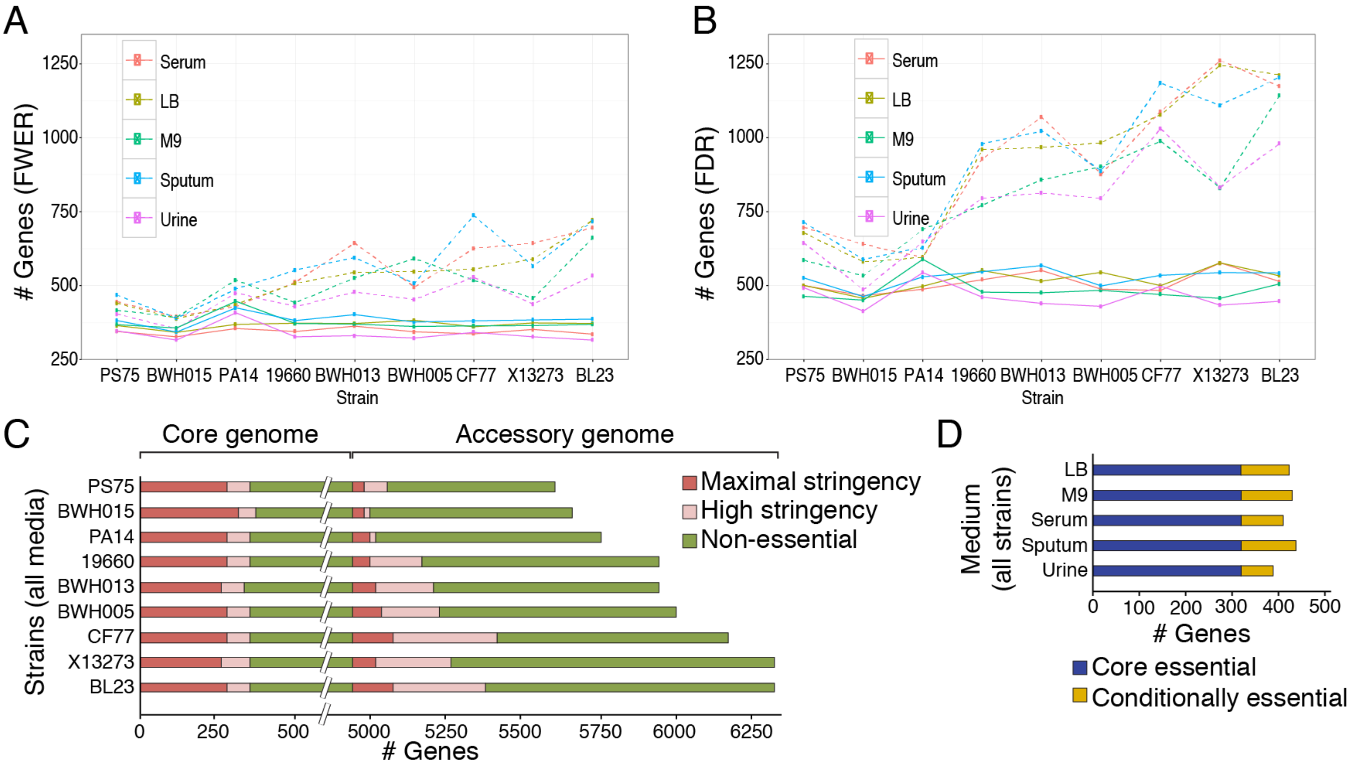


**Figure S5. Number of essential genes in genome as defined by *FiTnEss*.** A. maximal stringency (FWER) and B. high stringency (FDR) numbers of essential genes in whole genome for each strain and medium. Strains are ordered based on genome size, demonstrating the increase of accessory essential genes with genome size. Core genome essential genes are shown with solid lines, and total essential genes including the accessory genome essential genes are shown with dashed lines. C. Number of maximal stringency (red), high stringency (pink), and non-essential (green) genes common to each strain across all media, distributed between the 4903 core genes (left) and accessory genes (right). D. Number of essential genes common across all strains in each medium, highlighting the 321 core essential (blue) and conditionally essential (yellow) genes.

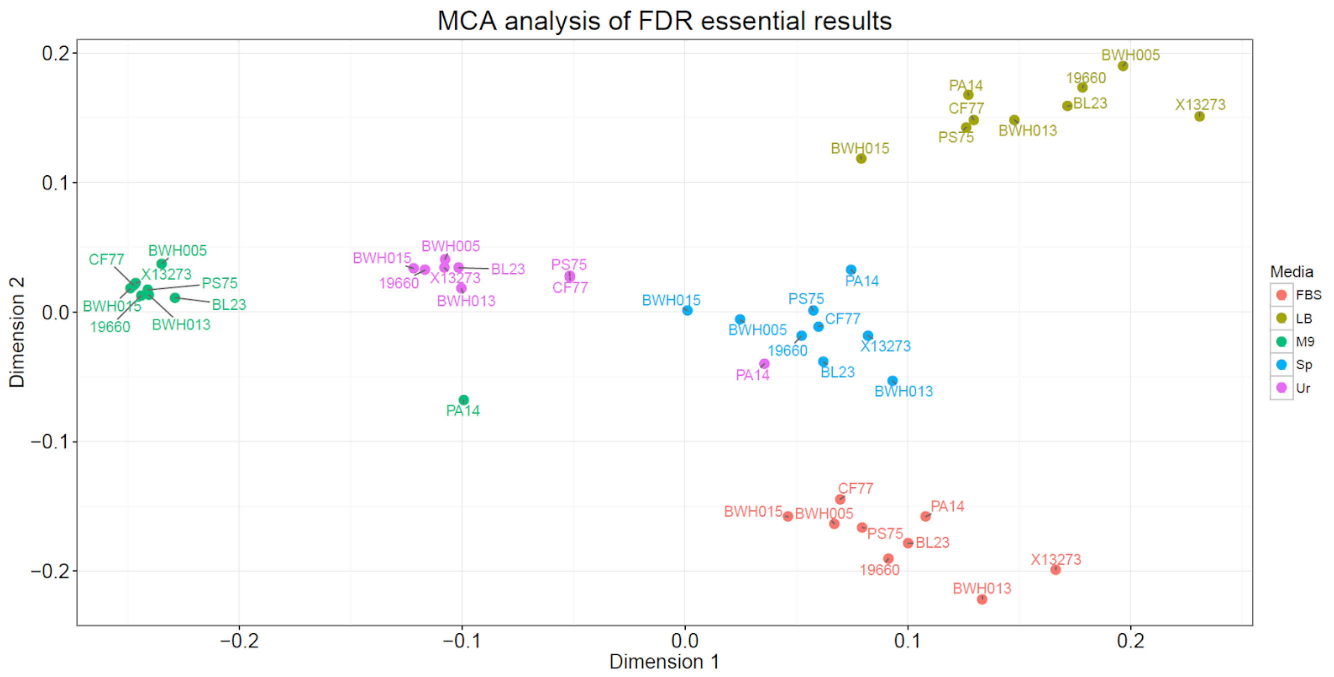


**Figure S6. MCA analysis of strain-media conditions.** MCA results of high-stringency essential genes identified by *FiTnEss* after FDR correction, projected on first two dimensions. Here we can see that conditions are grouped finely together based on their growth media, with PA14 being outlier in M9 minimal and urine media.

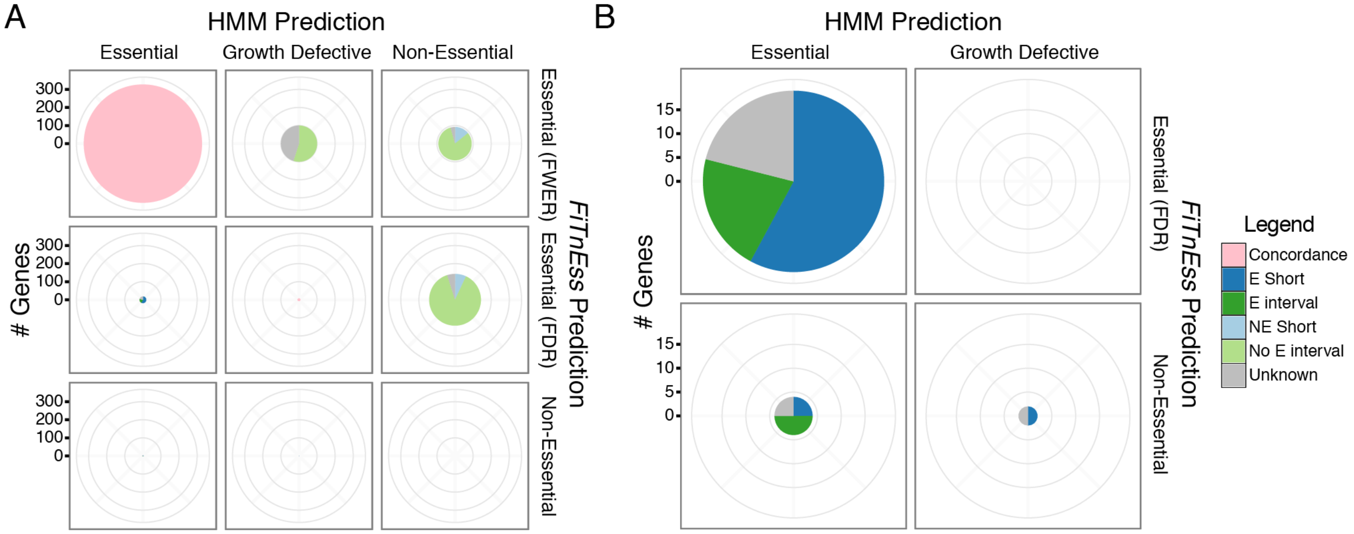


**Figure S7. Explanations of discrepancies between *FiTnEss* and TRANSIT (HMM method) predictions using the example PA14 on M9 media.** A. *FiTnEss* predictions of 'Essential (FWER)' (maximal stringency essential), 'Essential (FDR)' (high stringency essential) and 'Non-Essential' compared to HMM predictions of 'Essential', 'Growth Defective' and 'Non-Essential'. Pie sizes proportional to number of genes that fall into each category, with pie components indicating explanations of discrepant calls. Top left to bottom right diagonal panels are concordant calls colored in pink (data not shown for 'Non-Essential' vs 'Non-Essential' panel due to large size), thus the majority of all predictions are concordant between both methods. The 'Growth Defective' vs 'Essential (FWER)', 'Non-Essential' vs 'Essential (FWER)' and 'Non-Essential' vs 'Essential (FDR)' panels show genes that detected by *FiTnEss* as essential but are called less confidently (growth defective category) or completely non-essential by HMM. These genes for the most part fall unequivocally on the left mode of the ng distribution, and therefore justify *FiTnEss*'s calls. Closer examination of the data suggested that for most genes in this category, there are a moderately low number of reads that are distributed evenly across the genes. Since HMM focuses on consecutive runs of essential TA sites (that typically have near-zero reads), it has less power to detect the essentiality of such genes, whereas *FiTnEss* leverages all the TA sites in the gene and can thus more easily find a significant overall deficiency (**No E interval**). Another set of genes in this category are short genes with very few or no reads, flanked by non-essential genes with many reads. Here too the HMM's focus on runs of usually 5 or more essential TA sites reduces the method's power to detect essentiality (**NE Short**). B. Scaled-up versions of 'Essential' vs 'Essential (FDR)', 'Essential' vs 'Non-Essential', and 'Growth Defective' vs 'Non-Essential' are genes that are called essential or growth defective by HMM but are called less confidently or not essential at all by *FiTnEss*. Many of the genes in this category do not fall convincingly in the left mode of ng distribution. These genes tend to be short and seem to "mislead" HMM by being flanked by essential genes (**E short**). Finally, there is a small set of genes that HMM seems justified in called essential as they exhibit a clear interval of essential TA sites, typically an essential region of a large gene, but that interval is being "diluted" over the entire gene and thus reduces the power of *FiTnEss* to detect them (**E interval**). In summary, while there is complementarity between the methods, *FiTnEss* seems to be more powerful for calling essential genes than HMM, with only a very small tradeoff for genes where the latter has an advantage.

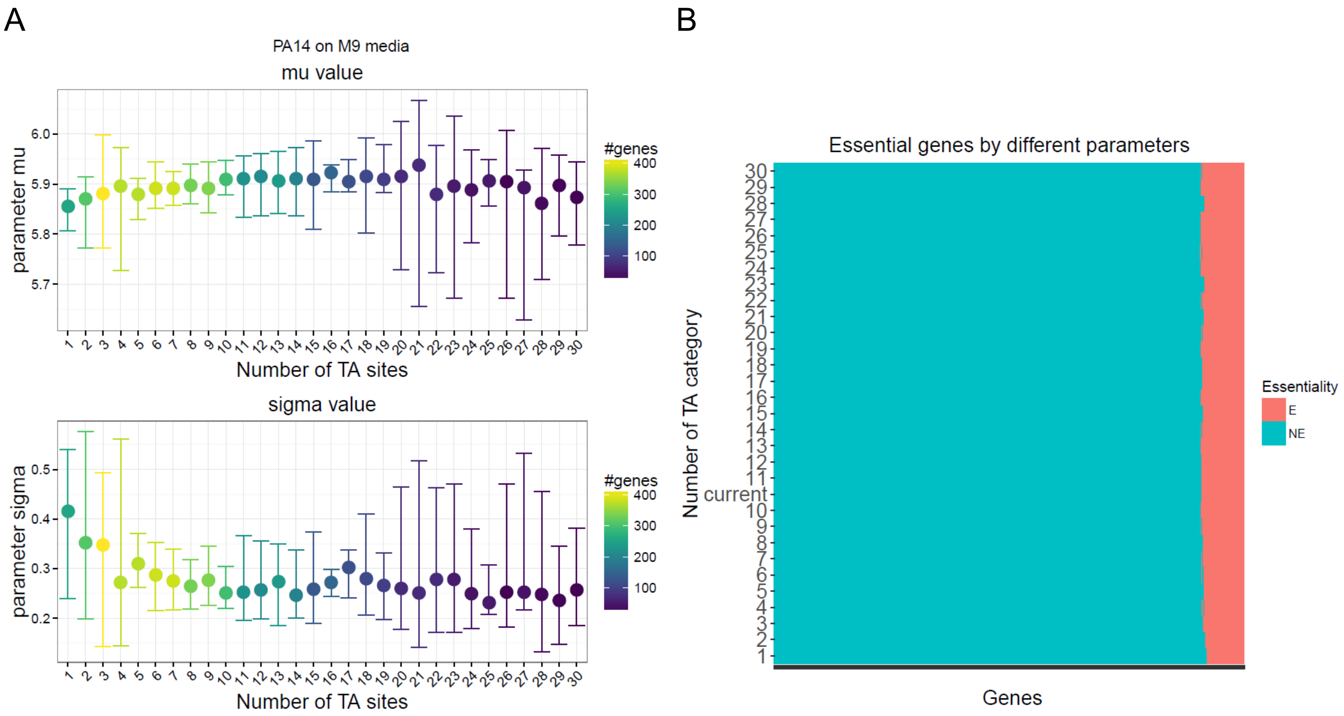


**Figure S8. Robustness test of *FiTnEss* parameters.** A. Variation of fitted parameters using genes with different number of TA site categories. Genes with $\boldsymbol{N_{TA}}$ number of TA sites ($\boldsymbol{N_{TA}} \in [\mathbf{1, 30}]$) were used to fit parameters respectively, and we repeated this process for 10 times in order to learn the variance. In this plot, dot showing the optimized parameter obtained from 10 runs for each $\boldsymbol{N_{TA}}$ category, while error bars showing 3 standard deviations around mean of those 10 runs. Color shows number of genes in each $\boldsymbol{N_{TA}}$ category. We found that parameters obtained using genes with 5-15 TA sites (median $\boldsymbol{N_{TA}} = \mathbf{9}$) were relatively consistent with small variations. In our analysis, we chose genes with 10 TA sites to fit parameters. B. Consistency of essential genes called by different parameters. We repeated essential gene calling process using optimized parameters obtained using genes of each $\boldsymbol{N_{TA}}$ category. Each row shows results using parameters obtained from corresponding $\boldsymbol{N_{TA}}$ categories shown on y-axis, with gene colored red to be essential, and blue to be non-essential. We could see that results are robust regardless of which $\boldsymbol{N_{TA}}$ category we used to fit parameters. Row with "current" label are our current results using genes with 10 TA sites.

### Tables

**Table S1. Summary of removed and usable TA insertion sites from analysis.**

| Strain | Total Genomic | Non-permissive | Homologous surrounding sequence | 5' and 3' gene end | Total Removed | Total Usable |
|---|---|---|---|---|---|---|
| 19660 | 83542 | 6558 | 1942 | 10274 | 17684 | 65858 |
| BL23 | 92065 | 6874 | 2720 | 11413 | 19732 | 72333 |
| BWH005 | 84887 | 6574 | 1194 | 10400 | 17196 | 67691 |
| BWH013 | 84102 | 6647 | 966 | 10120 | 16864 | 67238 |
| BWH015 | 78683 | 6374 | 786 | 9448 | 15785 | 62898 |
| CF77 | 89916 | 6936 | 2475 | 10826 | 19166 | 70750 |
| PA14 | 81328 | 6367 | 1122 | 9829 | 16499 | 64829 |
| PS75 | 75854 | 6199 | 671 | 9249 | 15309 | 60545 |
| X13273 | 88610 | 6793 | 1416 | 10849 | 18085 | 70525 |

**Table S2. Summary of TRANSIT (HMM) performance based on gene deletion growth profiles from Fig. S4.**

| | TRANSIT - HMM Prediction[a] | | |
|---|---|---|---|
| Growth category | Essential | Growth-defective | Non-Essential |
| Essential | 100% (15) | 80% (12) | 8% (7) |
| Intermediate | 0% (0) | 20% (3) | 14% (12) |
| Non-essential | 0% (0) | 0% (0) | 78% (66) |

[a]Strain-medium instances are in parentheses
